# Spatial artefact detection improves reproducibility of drug screening experiments

**DOI:** 10.1101/2025.04.07.647506

**Authors:** Aleksandr Ianevski, Kristen Nader, Swapnil Potdar, Alexandra Gorbonos, Filipp Ianevski, Ziaurrehman Tanoli, Jani Saarela, Tero Aittokallio

**Affiliations:** Institute for Molecular Medicine Finland (FIMM), HiLIFE, University of Helsinki, Helsinki, Finland; iCAN Digital Precision Cancer Medicine Flagship, University of Helsinki and Helsinki University Hospital, Helsinki, Finland; Institute for Cancer Research, Department of Cancer Genetics, Oslo University Hospital, Oslo, Norway; Oslo Centre for Biostatistics and Epidemiology (OCBE), Faculty of Medicine, University of Oslo, Oslo, Norway

**Author notes:** Correspondence: Aleksandr Ianevski,; Tero Aittokallio. These authors contributed equally to this work (Aleksandr Ianevski, Kristen Nader, Swapnil Potdar).

## Abstract

Reliable and reproducible drug screening experiments are essential for drug discovery and personalized medicine. Here, we demonstrate how systematic experimental errors negatively impact reproducibility, and that conventional quality control (QC) methods based on plate controls fail to detect these errors. To address this limitation, we developed a control-independent QC approach using normalized residual fit error (NRFE) to identify systematic errors in drug screening experiments. Comprehensive analysis of >100,000 duplicate measurements from the PRISM pharmacogenomic study revealed that the NRFE-flagged experiments show three-fold lower reproducibility between technical replicates. By integrating NRFE with existing QC methods to analyze 41,762 matched drug-cell line pairs between two datasets from the Genomics of Drug Sensitivity in Cancer project, we improved the cross-dataset correlation from 0.66 to 0.76. Available as an R package at https://github.com/IanevskiAleksandr/plateQC, plateQC provides a robust toolset for enhancing drug screening data reliability and consistency for basic research and translational applications.

## Introduction

High-throughput screening (HTS) of cancer cell lines and patient-derived samples has become central to drug discovery and personalized medicine. Large-scale pharmacogenomic initiatives, such as Cancer Cell Line Encyclopedia (CCLE)^1,2^, Genomics of Drug Sensitivity in Cancer (GDSC)^3,4^, and Profiling Relative Inhibition Simultaneously in Mixtures (PRISM)^5,6^, have expanded our understanding of drug responses in diverse genetic backgrounds. Despite their broad impact, several studies have reported problems regarding inter-laboratory consistency and inter-replicate reproducibility of drug response measurements^7–11^. These observations have led to valuable discussions about assay optimization strategies and best practices for cell-based drug response profiling, emphasizing the importance of robust validation approaches before translating preclinical findings.

Among various factors affecting drug screening data reproducibility^12^, one important consideration is the experimental data quality. Quality control in HTS drug experiments traditionally relies on simple plate control-based metrics and universal cutoff values, adhering to widely accepted standards such as Z-prime/Z-factor (Z’ > 0.5)^13^, Strictly Standardized Mean Difference (SSMD > 2)^5,14^ and signal-to-background ratio (S/B > 5)^13,15^ as the primary plate quality indicators even in the recent HTS studies ^4,16–20^. While these QC approaches have provided valuable and straightforward quality assessment for decades of HTS, they are fundamentally limited in their ability to detect many critical issues that can compromise experimental data quality. Control wells, by their nature, can only assess a fraction of the plate spatial area, and therefore the control-based QC metrics cannot capture systematic errors that affect the drug wells^21^.

The inherent challenges in HTS quality assessment stem from multiple factors. First, compound-specific issues - such as drug precipitation, stability changes during storage or assay conditions, carryover between wells during liquid handling, or interference with assay readouts – can significantly impact data quality, even when control wells appear adequate. Second, plate-specific artifacts - including evaporation gradients, systematic errors in multi-channel pipetting, and temperature-induced drift - can create spatial patterns of variability that may affect control and sample wells differently or occur in regions not covered by the controls. Third, position-dependent effects - such as striping or edge-well evaporation that leads to artificially high drug concentrations, or location-specific aggregation - introduce systematic errors that the control-based metrics fail to detect.

These quality issues often remain undetected when using the existing QC approaches and can therefore significantly impact the screening readouts and downstream analyses. Even plates passing traditional control-based quality metrics may still harbor systematic spatial errors that affect drug response measurements and dose-response curve fitting. These errors can lead to unreliable drug response quantifications using response metrics such as AUC or IC50 - particularly when spatial artifacts coincide with compound concentration patterns. In large-scale screens, such issues may cause inconsistent results between replicates, compromise the identification of true hits, and ultimately misdirect follow-up studies. These inconsistencies have direct consequences on the consistency of preclinical drug profiling results across different laboratories^22–26^.

To address these limitations, we developed a novel quality assessment approach based on normalized residual fit error (NRFE), which evaluates plate quality directly from drug-treated wells rather than relying on control wells. By analyzing deviations between the observed and fitted response values, while accounting for the variance structure of dose-response data, the method identifies systematic errors in the drug wells that the control-based metrics fail to detect. Analysis of four large-scale pharmacogenomic datasets (GDSC1, GDSC2, PRISM, and FIMM) demonstrates that the NRFE metric complements traditional control-based approaches; while control-based metrics excel at detecting assay-wide technical issues, NRFE captures drug-specific and position-dependent spatial artifacts. The integration of these orthogonal approaches therefore substantially improves our ability to identify unreliable drug response data, and enhance the quality, reproducibility and consistency of drug screening experiments.

## Normalized Residual Fit Error detects systematic plate artefacts

To systematically evaluate experimental plate quality in HTS experiments, we first examined traditional control-based metrics and identified their limitations. While metrics like Z-prime (Z′), SSMD, and signal-to-background ratio (S/B) have been widely adopted as industry standards, they rely solely on control wells to assess plate quality. Z-prime evaluates separation between positive and negative controls using means and standard deviations, SSMD quantifies the normalized difference between controls, and S/B measures the ratio of mean control signals (**Fig. 1a**, right top panel). Recognizing the need to detect artifacts in drug-containing wells, we developed the Normalized Residual Fit Error (NRFE) metric, which is based on deviations between the observed and fitted values in dose-response curves across all compound wells, and applies a binomial scaling factor to account for response-dependent variance (see Methods, **Fig. 1a**, right bottom panel).

**Figure 1.**
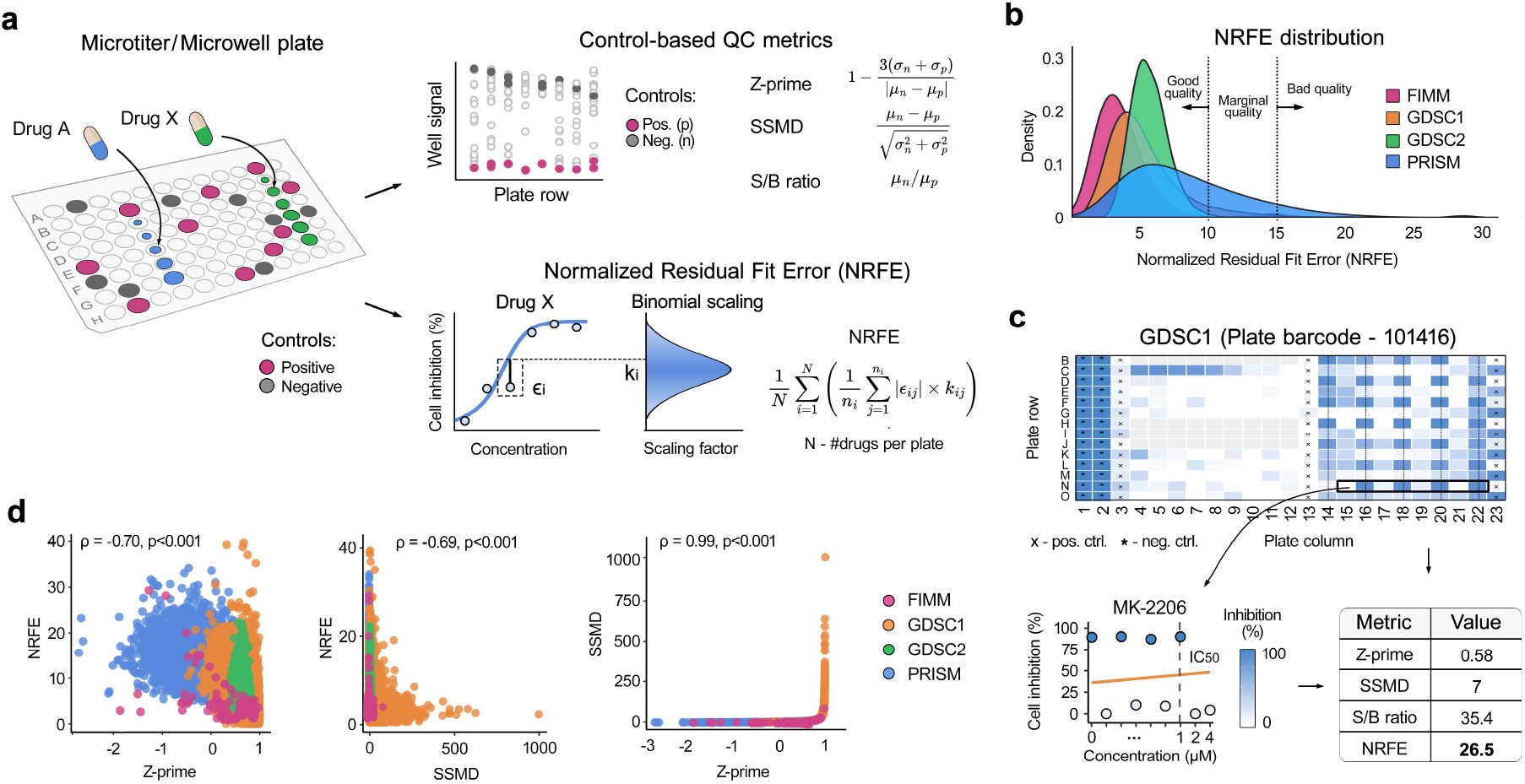
NRFE provides complementary quality assessment to traditional control-based metrics. (**a**) Calculation principles for drug screening plate quality control (QC) metrics: traditional control-based metrics (Z-prime, SSMD, S/B) are based on within plate positive and negative control wells (top), and NRFE approach makes use of dose-response curves from compound-containing wells (bottom). See Materials and Methods for the mathematical definitions of the metrics. (**b**) NRFE distributions across large-scale pharmacogenomic screens show relatively consistent quality baselines, with modal values of 2.8 (FIMM, n=1,044 plates), 3.9 (GDSC1, n=17,961), 5.1 (GDSC2, n=11,440), and 5.9 (PRISM, n=49,545). (**c**) Top panel: an example heatmap of the plate 101416 from the GDSC1 dataset shows column-wise striping artifacts on the right side of the plate. Bottom panels: an example of a compromised dose-response curve for compound MK-2206 (left) from plate 101416 that passed traditional quality metrics but was flagged by its extremely high NRFE value (right). (**d**) Correlation analysis between the quality metrics revealed a strong Spearman correlation between Z-prime and SSMD, but only a moderate negative correlation with NRFE (p < 0.001 for all comparisons, permutation test). Each point corresponds to a separate plate in the datasets.

To establish robust quality control thresholds for NRFE, we analyzed its distribution across 79,990 drug plates from four large-scale pharmacogenomic datasets (GDSC1, GDSC2, PRISM, and FIMM; **Fig. 1b, Suppl. Fig. 1**). These datasets were selected as they provide both dose-response measurements and plate location information required for systematic quality assessment. While GDSC1, GDSC2, and FIMM showed similar distributions, PRISM exhibited systematically higher NRFE values, likely due to its distinct experimental setup that uses pooled-cell screening format, longer treatment time (120h vs 72h), and luminex-based readout rather than standard cell viability assays. Notably, PRISM also showed substantially worse performance in the traditional control-based metrics (Z-prime and SSMD; **Suppl. Fig. 1**).

In the better-quality datasets, values exceeding three standard deviations corresponded to NRFE thresholds of approximately 10 in GDSC1 and 15 in both GDSC2 and FIMM datasets (**Fig. 1b**). These statistically-derived thresholds were validated using previously identified low-quality plates from internal FIMM screening data (see **Methods**), which led to NRFE values predominantly above 15 (**Suppl. Fig. 1a**). Based on this convergence of statistical analysis and internal validation, we defined three quality tiers: NRFE > 15 indicating low quality and requiring exclusion or careful review, 10-15 indicating borderline quality and requiring additional scrutiny, and NRFE < 10 indicating acceptable quality. These empirically validated NRFE thresholds complement the established control-based cut-offs, Z-prime (>0.5) and SSMD (>2).

The practical utility of NRFE became evident in its ability to identify problematic plates that passed the control-based quality control. For instance, plate 101416 from the GDSC1 dataset exhibited pronounced column-wise striping in the right half of the plate (**Fig. 1c**). This spatial pattern, likely arising from liquid handling irregularities, severely affected the dose-response relationships of multiple compounds, including MK-2206, which showed irregular jumpy dose responses that deviate from the expected sigmoid behavior (**Fig. 1c**, bottom panel). Despite these clear artifacts, traditional metrics indicated an acceptable quality (Z-prime = 0.58, SSMD = 7, S/B = 35.4), while an extremely high NRFE of 26.5 flagged the systematic quality issues (**Fig. 1c**, right-bottom panel). Additional examples of NRFE detecting systematic artifacts missed by the control-based metrics are shown in **Suppl. Fig. 2**.

Interestingly, when examining correlations between the QC metrics in these datasets, we found that S/B, which relies solely on averaged control signals without considering their variability, showed the weakest correlations with the other metrics (|ρ|<0.2, p<0.001, **Suppl. Fig. 3**). In contrast, Z-prime and SSMD were highly correlated (ρ = 0.99, p<0.001), indicating that they capture similar quality aspects, while NRFE showed only a moderate negative correlation with both of these metrics (Z-prime: ρ = -0.70, SSMD: ρ = -0.69, p<0.001) (**Fig. 1d**). This lower yet significant correlation indicates that NRFE considers distinct quality aspects, compared to the control-based metrics, supporting its use as a complementary approach for detecting systematic errors in drug-response assays.

### NRFE predicts the technical reproducibility of drug response measurements

We next examined whether plates with elevated NRFE levels exhibit reduced reproducibility in drug response measurements. For this analysis, we focused on the PRISM dataset, which, despite showing lower quality metrics overall, provided an ideal testbed with over 500,000 drug-cell line combinations tested across multiple plates. Within this extensive dataset, we identified 151,629 drug-cell line pairs with independent measurements on exactly two unique plates (**Fig. 2a**). To ensure reliable dose-response curve fitting, we further subselected 110,327 cases where the drugs were tested across more than three concentrations.

**Figure 2.**
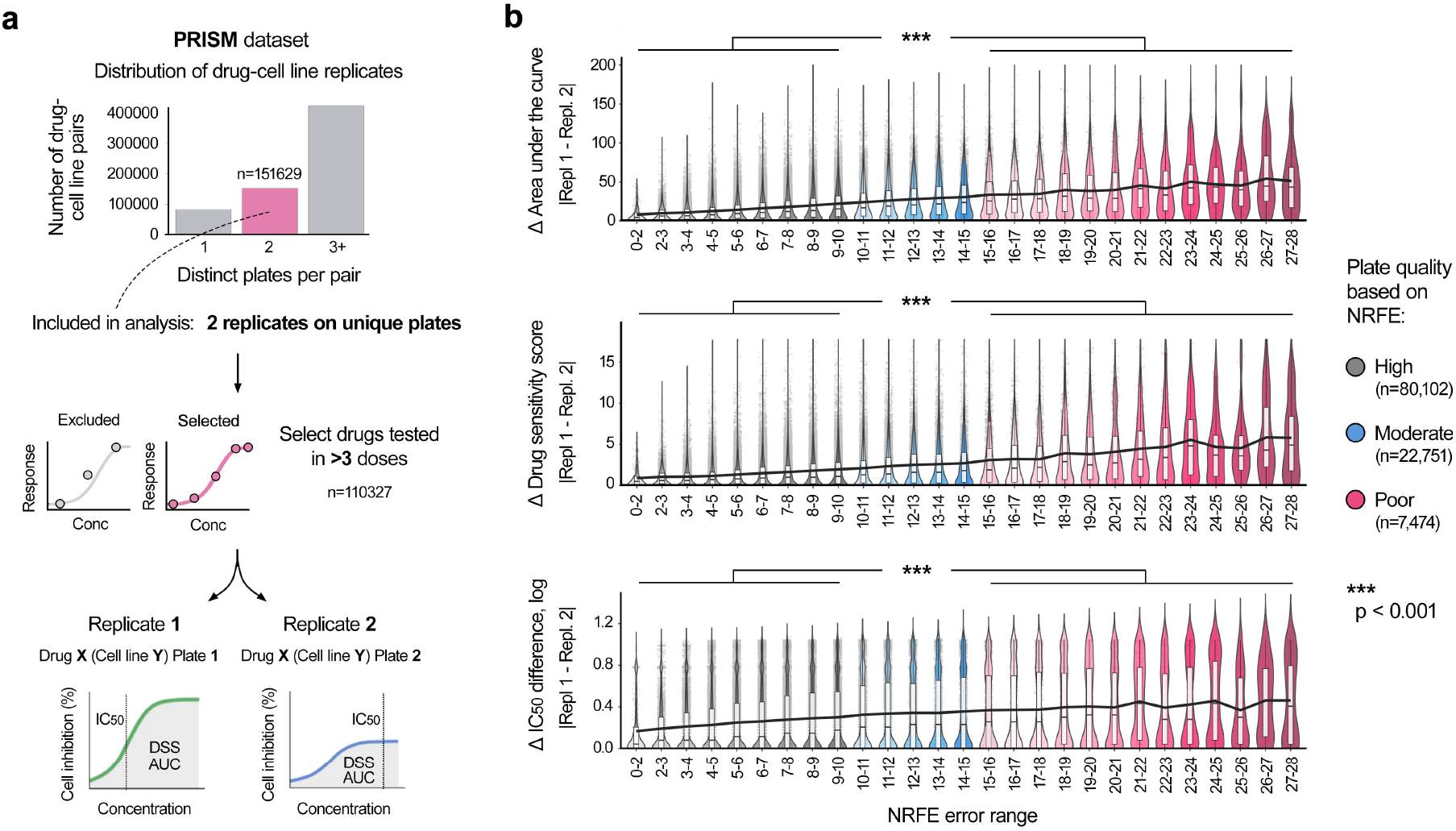
NRFE identifies drug response measurements with reduced technical reproducibility. (**a**) Overview of the PRISM dataset reproducibility analysis workflow: from more than 500,000 drug-cell line combinations, we identified 151,629 pairs with measurements on exactly two distinct plates, of which 110,327 had sufficiently many concentration points (>3 doses) for reliable curve fitting. (**b**) Impact of NRFE plate quality on the measurement reproducibility. Drug-cell line pairs were categorized by their plate NRFE values into high (gray, n=80,102), moderate (blue, n=22,751), and poor (red, n=7,474) quality categories. Black trend lines indicate the median values across NRFE categories, showing increasing variability between replicate measurements as a function of NRFE level for three drug response metrics: area under the curve (AUC), drug sensitivity score (DSS), and half maximal inhibitory concentration (IC_50_). The box plots show the median (central line), 25th and 75th percentiles (box edges), and the range within 1.5 times the interquartile range from the box (whiskers).

A striking pattern emerged when we categorized the drug-cell line measurements according to their plate NRFE values into three quality categories: high (NRFE<10, n=80,102), moderate (10≤NRFE≤15, n=22,751), and poor (NRFE>15, n=7,474), **Fig. 2b**. By comparing the replicate measurements between plates of different quality categories, we found that the pairs where at least one replicate came from a poor-quality plate showed substantially worse reproducibility, compared to the high-quality plates (p < 0.001, Wilcoxon test), with black trend lines emphasizing the consistent relationship between increasing NRFE values and measurement variability (**Fig. 2b**). This effect was consistent across three commonly-used drug response metrics: area under the curve (AUC, overall drug sensitivity), drug sensitivity score (DSS, normalized AUC)^27^, and half maximal inhibitory concentration (IC_50_, drug potency) values. For example, replicate measurements from high and moderate quality plates showed AUC differences of 11.8 on average, while those involving poor-quality plates exhibited average differences of 29.8 - a three-fold increase in response variability.

Notably, the poor-quality measurements comprised only 6.8% of the whole PRISM dataset, suggesting an opportunity to substantially improve data reliability through targeted removal of a small fraction of the most problematic plates. This finding has important implications for large-scale drug screening campaigns, which rely on maintaining broad coverage while ensuring sufficient data quality.

### Integration of quality control metrics improves cross-dataset consistency

After establishing the ability of NRFE to identify less reproducible measurements within the PRISM dataset, we next investigated whether combining multiple quality metrics could further improve consistency across different studies. For this analysis, we compared the drug response measurements between GDSC1 and GDSC2 datasets. From 84,657 matching drug-cell line pairs between the two datasets, we focused our analysis on 41,762 pairs that appeared exactly once in each dataset, enabling direct plate-level quality assessment (**Fig. 3a**). To quantify drug response consistently, we calculated drug sensitivity scores (DSS), a normalized area under the dose-response curve metric. Initial analysis of all matched pairs revealed a moderate cross-dataset correlation (Spearman ρ = 0.59 and Pearson r = 0.66; **Fig. 3a**, bottom panel), providing a baseline for evaluating quality-dependent drug response data consistency.

**Figure 3.**
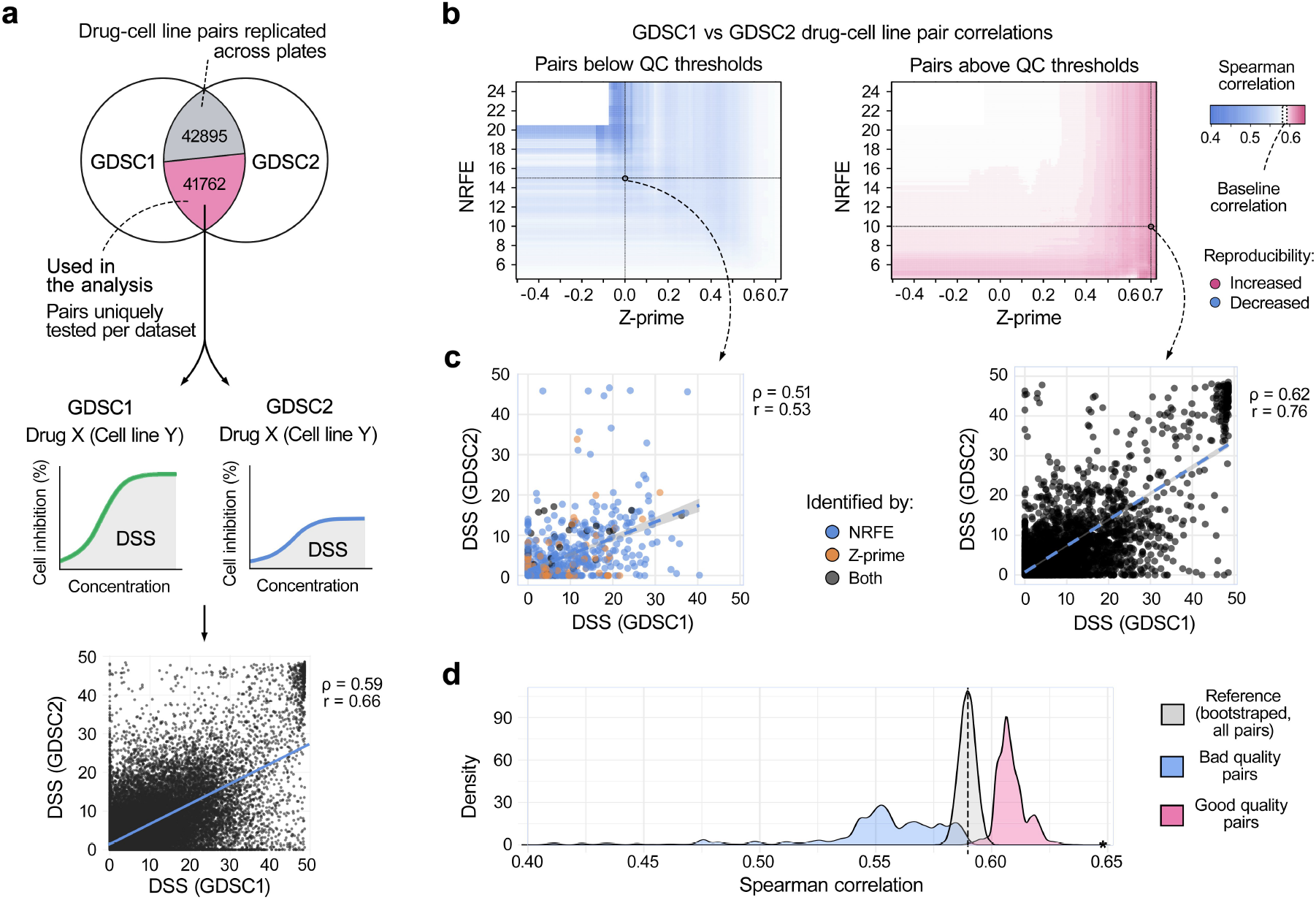
Quality control metrics contribute to drug response consistency across datasets. (**a**) Analysis workflow: Venn diagram of 84,657 matching drug-cell line pairs between GDSC1 and GDSC2 datasets, with 41,762 pairs appearing exactly once in each dataset, enabling direct plate-level quality assessment (top). Baseline correlation of drug sensitivity scores (DSS) between datasets shows moderate consistency (Spearman ρ = 0.59, Pearson r = 0.66) (bottom). (**b**) Impact of quality thresholds on consistency shown as correlation heatmaps across Z-prime (−0.5 to 0.75) and NRFE (4 to 25) ranges. Left: correlations for the pairs with poor quality metrics (Z-prime below or NRFE above threshold in either dataset), with dotted lines indicating quality thresholds (NRFE = 15, Z-prime = 0). Right: correlations for the pairs with good quality metrics (Z-prime above and NRFE below threshold in both datasets), with dotted lines marking quality thresholds (NRFE = 10, Z-prime = 0.7). White indicates the baseline correlation (ρ = 0.59), with red and blue showing increased and decreased consistency, respectively. (**c**) Impact of combined quality metrics on consistency: the pairs from poor-quality plates (NRFE > 15, Z-prime < 0) show reduced correlation (ρ = 0.51, r = 0.53), while the pairs from high-quality plates (Z-prime > 0.7, NRFE < 10) show improved correlation (ρ = 0.62, r = 0.76), compared to the baseline (ρ = 0.59, r = 0.66). (**d**) Statistical validation using bootstrap sampling to establish a reference baseline distribution, compared with distributions from poor-quality plates (Z-prime < 0 or NRFE > 15) and high-quality plates (Z-prime > 0.5 and NRFE < 10). We systematically explored various quality thresholds by calculating correlations at 0.02 intervals across the full range of Z-prime (−0.5 to 0.75) and NRFE (4 to 25) values. The asterisk on x-axis indicates top 2% highest-quality plates (NRFE < 6.36, Z-prime > 0.73, ρ = 0.65, r = 0.80). All differences significant at p < 0.001 (Kolmogorov-Smirnov test).

To systematically evaluate the combined impact of different quality metrics on the consistency, we stratified the matched drug-cell line pairs based on both their plate Z-prime (range -0.5 to 0.75) and NRFE values (range 4 to 25). For each threshold combination, we calculated Spearman correlations separately for the drug-cell line pairs with poor quality control metrics (Z-prime below the threshold or NRFE above the threshold in either dataset) and good quality control metrics (Z-prime above the threshold and NRFE below the threshold in both datasets). Visualization of these correlations as heatmaps demonstrated that the drug-cell line pairs derived from plates with poor quality metrics showed decreased consistency, while pairs with acceptable quality metrics exhibited improved consistency, when compared to the baseline correlation of ρ=0.59 (**Fig. 3b**).

The added value of combining the quality metrics became evident when examining specific threshold combinations across the dataset of 41,762 drug-cell line pairs (**Fig. 3c**). For example, drug-cell line pairs from the poor-quality plates with both high NRFE and low Z-prime values (NRFE > 15 and Z-prime < 0) showed a reduced cross-dataset correlation (ρ=0.51, r=0.53), while pairs from the plates with excellent quality based on both metrics (Z-prime > 0.7 and NRFE < 10) demonstrated an increased correlation (ρ=0.62 and r=0.76), when comparing to the baseline correlation (ρ=0.59 and r=0.66). These improvements in drug response data consistency across large-scale pharmacogenomic datasets demonstrate how integrating multiple quality metrics can systematically identify more reliable drug response measurements.

To statistically validate these findings, we used a bootstrap sampling of the matched drug-cell line pairs to establish a reference distribution for the baseline consistency (**Fig. 3d**). Two quality groups were then compared against the reference distribution: lower-quality pairs (defined by either Z-prime < 0 or NRFE > 15) and high-quality pairs (defined by both Z-prime > 0.5 and NRFE < 10). Both groups showed significant differences compared to the reference distribution (p < 0.001, Kolmogorov-Smirnov test; **Fig. 3d**). Further analysis of more stringent thresholds revealed that the top 2% highest-quality plates (NRFE < 6.36 and Z-prime > 0.73) achieved even stronger consistency, with correlations of ρ=0.65 and r=0.80 (**Fig. 3d**, asterisk). These results demonstrate how combining traditional control-based metrics with NRFE provides a robust framework for identifying reliable drug response measurements across drug screening studies.

## Discussion

Preclinical drug response reproducibility remains one of the most critical challenges in drug development and personalized medicine studies, as non-reproducible preclinical findings compromise therapeutic success in follow-up studies and clinical trials, with oncology having the highest failure rate compared to other therapeutic areas^22,23^. Our comprehensive analysis reveals that this reproducibility challenge stems not only from technical variability in drug screening assays but also from fundamental limitations in the current quality assessment approaches. Traditional control-based metrics, while valuable for detecting assay-wide technical issues, are inherently limited by their focus on control wells, potentially missing systematic errors that affect the drug response measurements.

The NRFE metric overcomes these limitations by assessing systematic errors directly in drug-treated wells, hence providing a complementary approach to traditional quality control. Our analysis of the relationships between quality metrics revealed their distinct roles - while Z-prime and SSMD showed a strong correlation (ρ = 0.99) in detecting assay-wide issues, NRFE showed a moderate negative correlation with these metrics (ρ ≈ -0.63), indicating its ability to capture different quality aspects. As shown by the representative examples in **Fig. 3c**, these metrics identify distinct types of experimental artifacts, indicating their complementary roles in comprehensive quality assessment of drug response measurements. The weak correlations of signal-to-background ratio with the other metrics (|ρ| < 0.2) questions its continued use in screening protocols, particularly given that it ignores the control well variability.

NRFE metric offers several practical advantages for drug screening workflows. It requires only percent inhibition (or viability) values, enabling quality assessment even without raw plate reader data. However, since NRFE relies on dose-response curve fitting, it is specifically designed for multi-dose assays, highlighting the continued importance of control-based metrics for single-dose screens. To facilitate adoption of this integrated quality control approach, we developed the plateQC R package that implements both control-based and drug-based metrics with interactive visualizations. The R package provides comprehensive tools for plate-level quality assessment, including automated detection of spatial artifacts, interactive visualization of dose-response curves, and statistical summaries of quality metrics. We recommend examining plates that fall below the established quality thresholds (see **Supplementary Figure 2**).

Analysis of four large-scale pharmacogenomic datasets (GDSC1, GDSC2, PRISM, and FIMM) demonstrates the broad applicability of NRFE approach across diverse screening platforms, assays and protocols. Notably, integration of NRFE with traditional metrics, such as Z-prime, enabled the identification of more reliable measurements in both technical replicates and cross-dataset comparisons, hence further improving data reproducibility and consistency. Since these large-scale datasets underpin numerous preclinical discovery and computational modeling efforts, improved quality control is expected to enhance the reliability of the downstream analyses and reduce the costs throughout the drug development process. The improved reliability and translatability of the preclinical findings will accelerate the production of new and desperately-needed therapies for many diseases.

In conclusion, this work establishes a comprehensive framework for quality control in high-throughput drug screening by combining control-based and drug-specific error detection approaches. While multiple factors influence experimental reproducibility, systematic identification of reliable measurements represents an important step toward more robust preclinical data generation. These principles may extend beyond drug screening to other high-throughput technologies, such as CRISPR-Cas9 screens, ultimately contributing to more effective therapeutic development strategies.

## Materials and Methods

### Calculation of Quality Control Metrics

Traditional control-based metrics (Z-prime, SSMD, and S/B) are calculated using positive and negative control wells. The Z-prime factor (also known as Z′ or Z-factor), being the most widely adopted industry standard, quantifies the separation between positive (p) and negative (n) control wells, calculated as Z′ = 1 - (3σ_p_ + 3σ_n_)/|μ_p_ - μ_n_|, where σ and μ represent standard deviation and mean, respectively, over the plate controls. Strictly Standardized Mean Difference (SSMD) is computed as 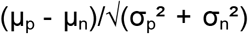. Signal-to-background ratio is calculated as S/B = μ_p_/μ_n_.

The Normalized Residual Fit Error (NRFE) is defined based on normalized residuals from a four-parameter log-logistic (LL4) model fit. We chose the widely-used LL4 model in the present study, but note that NRFE can also be calculated based on other dose-response functions. The LL4 model was defined as f(x) = d + (a - d)/(1 + (x/c)^b), where a is the minimum asymptote, d is the maximum asymptote, b is the slope, c is the inflection point (IC_50_), and x is the drug concentration. The residual errors ε_ij_ at each concentration point j was calculated as |y_ij_ - f(x_ij_)|, where y_ij_ is the observed response for drug i at concentration point j. NRFE is then computed as 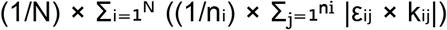, where N is the number of drug wells per plate, n_i_ is the number of concentrations for drug i, and k_ij_ is the binomial scaling factor. The scaling factor k_ij_ is calculated as 1 + (p × (1-p)/0.25), where p is the fitted response proportion (f(x_ij_)/100). This scaling factor adjusts for the weight of residuals based on their position in the dose-response curve, with the maximal weight at 50% response. Quality thresholds were established as NRFE > 15 for low quality requiring review, 10-15 for marginal quality needing scrutiny, and NRFE <10 for acceptable quality.

### Statistical assessment of correlation differences

To assess whether the correlations differed between quality-stratified groups, we first established a reference distribution of expected correlations by bootstrapping the original drug sensitivity scores (DSSs) (10,000 resamples with replacement). For each resample, we calculated the Spearman correlation between GDSC1 and GDSC2 dataset DSS scores, creating a distribution of background correlations under typical conditions. We then compared this reference distribution against the correlation distributions from two quality groups: low-quality plates (defined by NRFE > 15 or Z-prime < 0) and high-quality plates (NRFE < 10 and Z-prime > 0.5). Statistical significance was assessed using the two-sample Kolmogorov-Smirnov tests, which evaluates the null hypothesis that the two samples come from the same distribution. Effect sizes were calculated as standardized mean differences between each group and the reference distribution was, computed as the difference in means divided by the pooled standard deviation. All statistical analyses were performed in R v4.2.3 using the boot^28^ (v. 1.3-28.1) and stats packages.

## Data and Code Availability

The PRISM and GDSC (GDSC1 and GDSC2) datasets were accessed through DepMap Portal (https://depmap.org/portal/data_page/?tab=customDownloads). The FIMM dataset was accessed through PharmacoDB (https://pharmacodb.ca/) and extended with an internal dataset of 16 poor-quality plates that is available from the corresponding authors upon a reasonable request.

The plateQC R package implements all quality control metrics described in this study, along with interactive plate visualizations and robust outlier detection. The R package is freely available at https://github.com/IanevskiAleksandr/plateQC. Analysis scripts used to generate the results and figures are available from the corresponding authors upon reasonable request.

## Acknowledgements

We thank the DDCB core facility (FIMM HTB unit) supported by the University of Helsinki and Biocenter Finland. Funding support: AI: Ida Montin Foundation grant. TA: Research Council of Finland (grants 340141, 344698); the Cancer Society of Finland, the Norwegian Cancer Society (grants 216104 and 273810), the Sigrid Jusélius Foundation, and iCAN – Digital Precision Cancer Medicine Flagship (iCAN-MULTIDRUG).

## References

1. Barretina, J. et al. The Cancer Cell Line Encyclopedia enables predictive modelling of anticancer drug sensitivity. Nature 483, 603–607 (2012).

2. M, G. et al. Next-generation characterization of the Cancer Cell Line Encyclopedia. Nature 569, (2019).

3. Yang, W. et al. Genomics of Drug Sensitivity in Cancer (GDSC): a resource for therapeutic biomarker discovery in cancer cells. Nucleic Acids Res. 41, D955–D961 (2013).

4. Garnett, M. J. et al. Systematic identification of genomic markers of drug sensitivity in cancer cells. Nature 483, 570–575 (2012).

5. Corsello, S. M. et al. Discovering the anticancer potential of non-oncology drugs by systematic viability profiling. Nat. Cancer 1, 235–248 (2020).

6. Yu, C. et al. High-throughput identification of genotype-specific cancer vulnerabilities in mixtures of barcoded tumor cell lines. Nat. Biotechnol. 34, 419–423 (2016).

7. Weinstein, J. N. & Lorenzi, P. L. Discrepancies in drug sensitivity. Nature 504, 381–383 (2013).

8. Hatzis, C. et al. Enhancing Reproducibility in Cancer Drug Screening: How Do We Move Forward? Cancer Res. 74, 4016–4023 (2014).

9. Haverty, P. M. et al. Reproducible pharmacogenomic profiling of cancer cell line panels. Nature 533, 333–337 (2016).

10. Mpindi, J. P. et al. Consistency in drug response profiling. Nature 540, E5–E6 (2016).

11. Bouhaddou, M. et al. Drug response consistency in CCLE and CGP. Nature 540, E9–E10 (2016).

12. Niepel, M. et al. A Multi-center Study on the Reproducibility of Drug-Response Assays in Mammalian Cell Lines. Cell Syst. 9, 35-48.e5 (2019).

13. Zhang, J.-H., Chung, T. D. Y. & Oldenburg, K. R. A Simple Statistical Parameter for Use in Evaluation and Validation of High Throughput Screening Assays. SLAS Discov. 4, 67–73 (1999).

14. Zhang, X. D. A pair of new statistical parameters for quality control in RNA interference high-throughput screening assays. Genomics 89, 552–561 (2007).

15. Potdar, S. et al. Breeze 2.0: an interactive web-tool for visual analysis and comparison of drug response data. Nucleic Acids Res. 51, W57–W61 (2023).

16. Jaaks, P. et al. Effective drug combinations in breast, colon and pancreatic cancer cells. Nature 603, 166–173 (2022).

17. Ayuda-Durán, P. et al. Standardized assays to monitor drug sensitivity in hematologic cancers. Cell Death Discov. 9, 1–13 (2023).

18. Hermansen, J. U. et al. A tumor microenvironment model of chronic lymphocytic leukemia enables drug sensitivity testing to guide precision medicine. Cell Death Discov. 9, 1–10 (2023).

19. Acanda De La Rocha, A.M. et al. Feasibility of functional precision medicine for guiding treatment of relapsed or refractory pediatric cancers. Nat. Med. 30, 990–1000 (2024).

20. Hirt, C. K. et al. Drug screening and genome editing in human pancreatic cancer organoids identifies drug-gene interactions and candidates for off-label therapy. Cell Genomics 2, 100095 (2022).

21. Goktug, A. N. et al. Data Analysis Approaches in High Throughput Screening. In Drug Discovery (IntechOpen, 2013). doi:10.5772/52508.

22. Baker, M. 1,500 scientists lift the lid on reproducibility. Nature 533, 452–454 (2016).

23. Begley, C. G. & Ellis, L. M. Raise standards for preclinical cancer research. Nature 483, 531–533 (2012).

24. Honkala, A., Malhotra, S. V., Kummar, S. & Junttila, M. R. Harnessing the predictive power of preclinical models for oncology drug development. Nat. Rev. Drug Discov. 21, 99–114 (2022).

25. Haibe-Kains, B. et al. Inconsistency in large pharmacogenomic studies. Nature 504, 389–393 (2013).

26. Mullard, A. Preclinical cancer research suffers another reproducibility blow. Nat. Rev. Drug Discov. 21, 89–89 (2022).

27. Yadav, B. et al. Quantitative scoring of differential drug sensitivity for individually optimized anticancer therapies. Sci. Rep. 4, 5193 (2014).

28. Canty, A. & Ripley, B. boot: Bootstrap Functions (Originally by Angelo Canty for S). 1.3-31 10.32614/CRAN.package.boot (1999).

