## Supplementary Information for "Spatial artefact detection improves reproducibility of drug screening experiments"

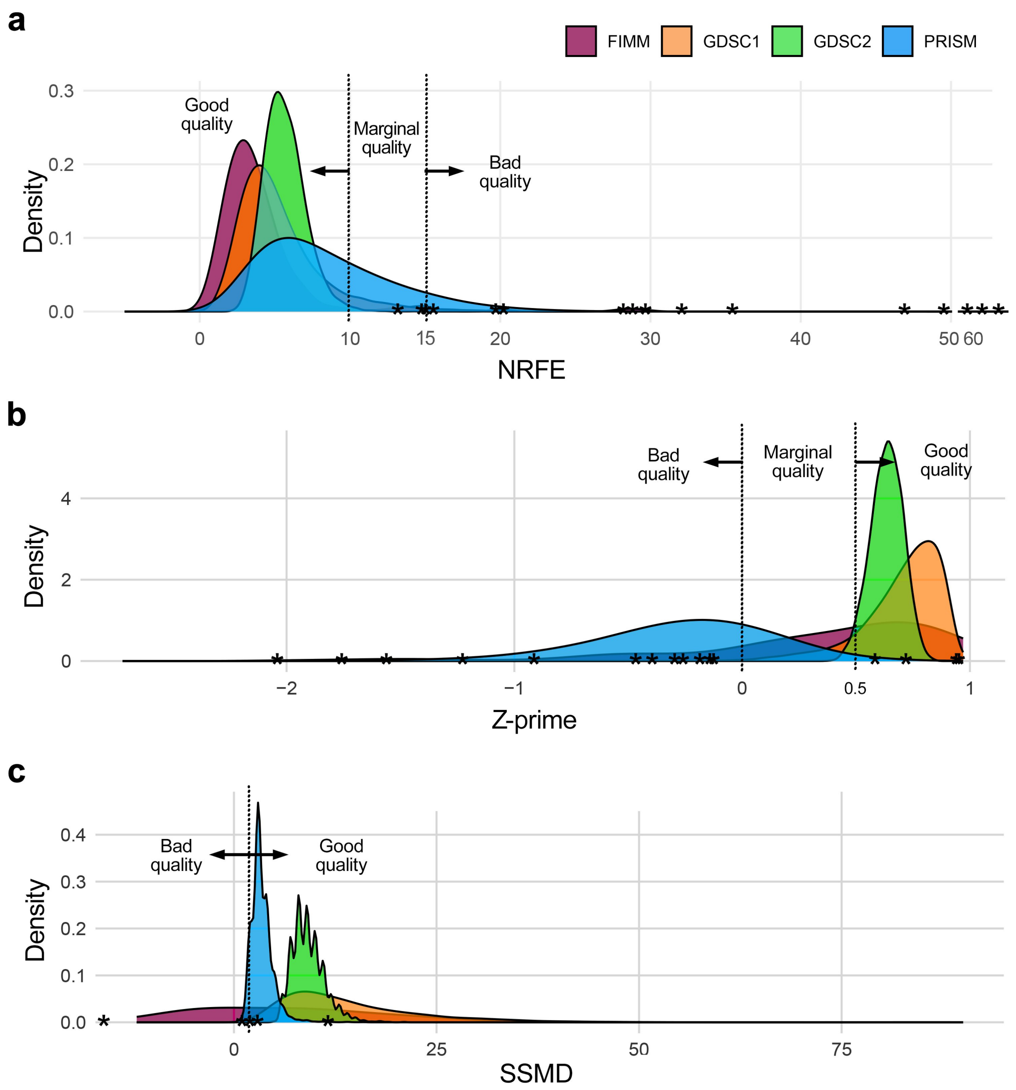


**Supplementary Figure 1 | Quality metric distributions across four pharmacogenomic screening datasets.** (**a**) Normalized Relative Fit Error (NRFE) distributions showing the quality thresholds: values >15 indicate poor quality plates requiring exclusion or detailed review, 10-15 indicate marginal quality needing additional scrutiny, and <10 represent acceptable quality plates. Stars indicate 16 manually curated poor-quality plates from the FIMM dataset that were visually identified to contain systematic artifacts. (**b**) Z-prime factor distributions demonstrating assay separation between positive and negative controls across all plates. Z-prime values >0.5 indicate excellent assay quality, 0-0.5 indicate marginal quality requiring additional scrutiny, and Z'<0 indicate poor separation between controls necessitating assay optimization or plate exclusion. (**c**) Strictly Standardized Mean Difference (SSMD) distributions measuring the effect size between control populations. SSMD >2 indicates good quality and SSMD <2 poor quality necessitating careful evaluation or exclusion.

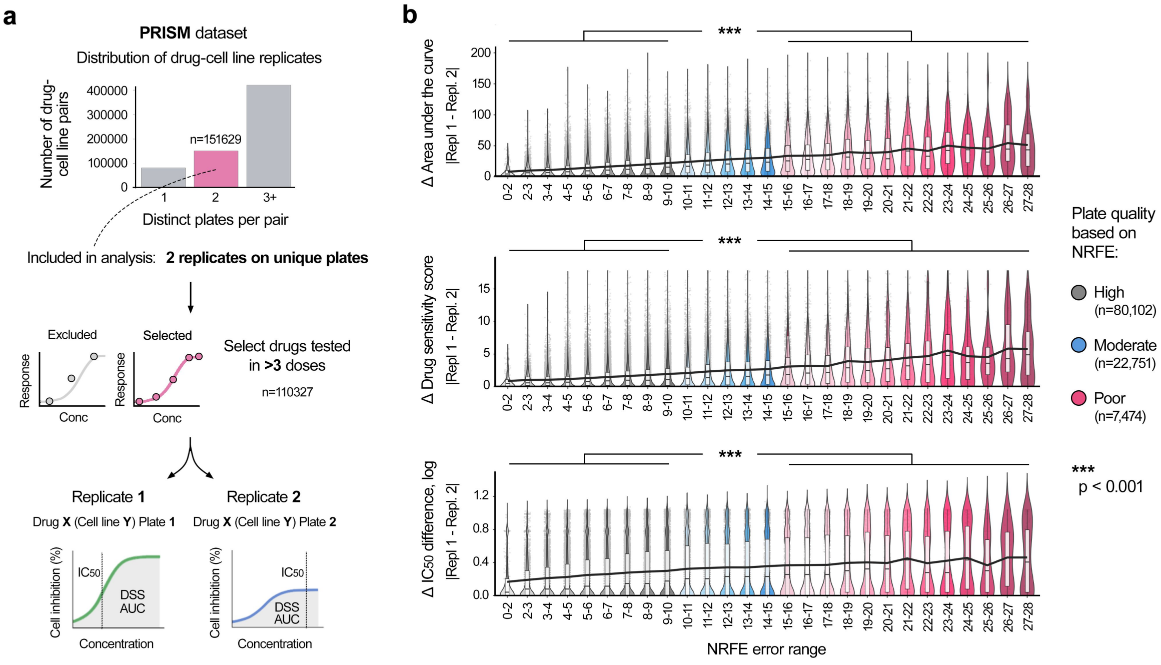


**Supplementary Figure 2 | NRFE identifies systematic artefacts in drug screening datasets not detected by the standard metrics.** (**a**) An example plate from GDSC1 dataset showing column-wise striping in the right half (left), which affects dose-response patterns of compounds with irregular dose-responses (middle). This systematic pattern was detected only by NRFE and Z-prime metrics (right, bold). (**b**) Another GDCS1 plate shows technical artefacts with potential precipitation/solubility issues (left), illustrated by sporadic drug response variations across concentrations of drug responses (middle), and identified solely by the NRFE metric (right). (**c**) An example of systematic dispensing artefacts in a drug plate from GDSC2 dataset (left), visible as abnormal variations specifically in the first and third drug doses across multiple compounds, which results in irregular jumpy behavior at these concentrations in the dose-response curves (middle), and was detected uniquely through the NRFE analysis (right). (**d**) An example of the plate from PRISM dataset demonstrates a checkerboard-like pattern of alternating high- and low-value responses (left), reflecting potential systematic dispensing or temperature gradient effects, visible as alternating intensity fluctuations in dose-response profiles (middle), and identified by the NRFE analysis and S/B ratio (right).

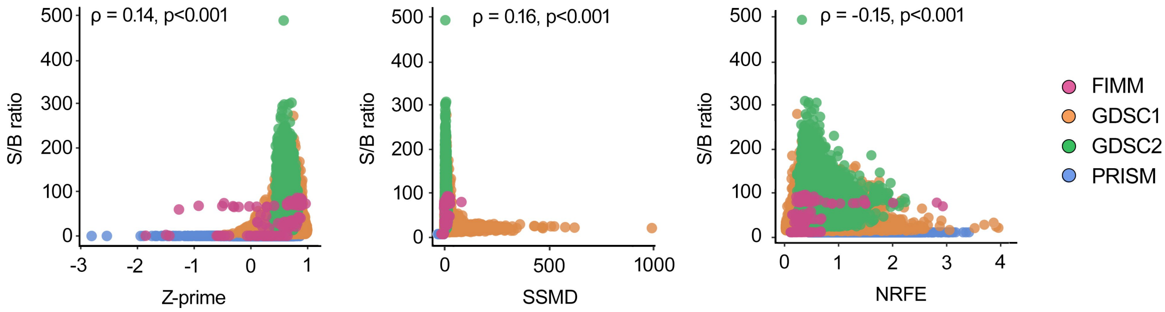


**Supplementary Figure 3 |** Correlation analysis between signal-to-background (S/B) ratio and the other quality metrics including Z-prime (left), SSMD (middle), and NRFE (right). Spearman correlations (ρ) are shown in each plot, with $p < 0.001$ for all comparisons, based on the permutation test.
